# Identification of plant enhancers and their constituent elements by STARR-seq in tobacco leaves

**DOI:** 10.1101/2020.02.17.952747

**Authors:** Tobias Jores, Jackson Tonnies, Michael W Dorrity, Josh T Cuperus, Stanley Fields, Christine Queitsch

## Abstract

Genetic engineering of *cis*-regulatory elements in crop plants is a promising strategy to ensure food security. However, such engineering is currently hindered by our limited knowledge of plant *cis*-regulatory elements. Here, we adapted STARR-seq — a technology for the high-throughput identification of enhancers — for its use in transiently transformed tobacco leaves. We demonstrate that the optimal placement in the reporter construct of enhancer sequences from a plant virus, pea and wheat was just upstream of a minimal promoter, and that none of these four known enhancers was active in the 3′-UTR of the reporter gene. The optimized assay sensitively identified small DNA regions containing each of the four enhancers, including two whose activity was stimulated by light. Furthermore, we coupled the assay to saturation mutagenesis to pinpoint functional regions within an enhancer, which we recombined to create synthetic enhancers. Our results describe an approach to define enhancer properties that can be performed in potentially any plant species or tissue transformable by *Agrobacterium* and that can use regulatory DNA derived from any plant genome.

**One-sentence summary:** We developed a high-throughput assay in transiently transformed tobacco leaves that can identify enhancers, characterize their functional elements and detect condition-specific enhancer activity.

## INTRODUCTION

In a time of climate change and increasing human population, crop plants with higher yields and improved response to abiotic stresses will be required to ensure food security. As many of the beneficial traits in domesticated crops are caused by mutations in *cis*-regulatory elements, especially enhancers, genetic engineering of such elements is a promising strategy for improving crops (Swinnen et al., 2016; Scheben et al., 2017). However, this strategy is currently not feasible at large scale due to our limited knowledge of *cis*-regulatory elements in plants.

As in animals, plant gene expression is controlled by *cis*-regulatory elements such as minimal promoters and enhancers. A minimal promoter is the DNA sequence necessary and sufficient to define a transcription start site and recruit the basal transcription machinery. Such minimal promoters generally lead to low levels of expression (Andersson and Sandelin, 2020). Enhancers are DNA sequences that can increase the basal transcription level established by minimal promoters. Enhancers serve as binding sites for transcription factors that interact with the basal transcription machinery to increase its rate of recruitment, transcription initiation, and/or elongation (Weber et al., 2016; Marand et al., 2017; Andersson and Sandelin, 2020). In contrast to promoters, enhancers function independently of their orientation. They can occur upstream or downstream of the minimal promoter and are active over a wide range of distances (Banerji et al., 1981, 1983; Chandrasekharappa and Subramanian, 1987). Enhancers can interact with minimal promoters that are several kilobases away, with such long-distance interactions assembled by chromatin loops that bring the enhancer and minimal promoter into close proximity (Amano et al., 2009; Studer et al., 2011; Weber et al., 2016; Ricci et al., 2019).

Enhancers can be identified by self-transcribing active regulatory region sequencing (STARR-seq), a massively parallel reporter assay (Arnold et al., 2013). Here, candidate enhancer sequences are inserted into the 3’ untranslated region (3′-UTR) of a reporter gene under the control of a minimal promoter. If an insert has enhancer activity, it can upregulate its own transcription. The resulting transcript can be detected by next generation sequencing and linked to its corresponding enhancer element, which is incorporated within the mRNA. This method has been widely used in *Drosophila* and human cells (Arnold et al., 2013, 2014; Liu et al., 2017a, 2017b; Wang et al., 2018). In plants, STARR-seq has been described in only two studies that applied the method to the monocot species rice and maize (Sun et al., 2019; Ricci et al., 2019). These studies analyzed the enhancer activity of fragments in large genomic libraries obtained from sheared rice DNA (Sun et al., 2019) or from transposase-digested maize DNA (Ricci et al., 2019). The latter approach enriches the library for fragments from open chromatin regions of the genome where active enhancers reside (Buenrostro et al., 2013; Andersson and Sandelin, 2020). Both these previous plant STARR-seq studies relied on species-specific protoplasts as recipient cells for the assay. However, efficient protoplasting and transformation protocols have been established for only a few species. Furthermore, protoplasts are often fragile and might not respond to external stimuli in the same way as intact plants.

Here, we established a STARR-seq assay that uses transient expression of STARR-seq libraries in tobacco leaves. This assay bypasses the need for a species-specific protoplasting protocol and instead relies on efficient *Agrobacterium*-mediated transformation. Among species that are amenable to transformation with *Agrobacteria*, tobacco combines fast and robust growth with convenient transformation by syringe-infiltration of intact leaves. As transcription factors are highly conserved among plant species (Lehti-Shiu et al., 2017; Wilhelmsson et al., 2017), the versatile tobacco system can serve as a proxy for many plant species, including crops. We optimized the placement of the enhancer candidates to provide an optimal dynamic range and performed proof-of-principle experiments to demonstrate that the assay can detect enhancers and characterize the underlying functional elements. Furthermore, we show that our *in planta* assay is capable of detecting light-dependent changes of the transcriptional activity of known light-sensitive enhancers.

## RESULTS

### The positioning of enhancers strongly impacts their activity in tobacco STARR-seq

Transient expression in tobacco leaves is a well-established method for reporter assays. We tested whether STARR-seq, a massively parallel reporter assay to identify active *cis*-regulatory elements, could be performed by transient expression of libraries in tobacco. We created a reporter construct with a green fluorescent protein (GFP) gene under control of the Cauliflower mosaic virus 35S minimal promoter and the 35S core enhancer (Fang et al., 1989; Benfey et al., 1990) (subdomains A1 and B1-3). *Agrobacterium tumefaciens* cells harboring this construct were used to transiently transform leaves of 3-4 week old tobacco (*Nicotiana benthamiana*) plants. After two days, the resulting mRNAs were extracted from the transformed leaves and analyzed by next generation sequencing (Figure 1A).

**Figure 1.**
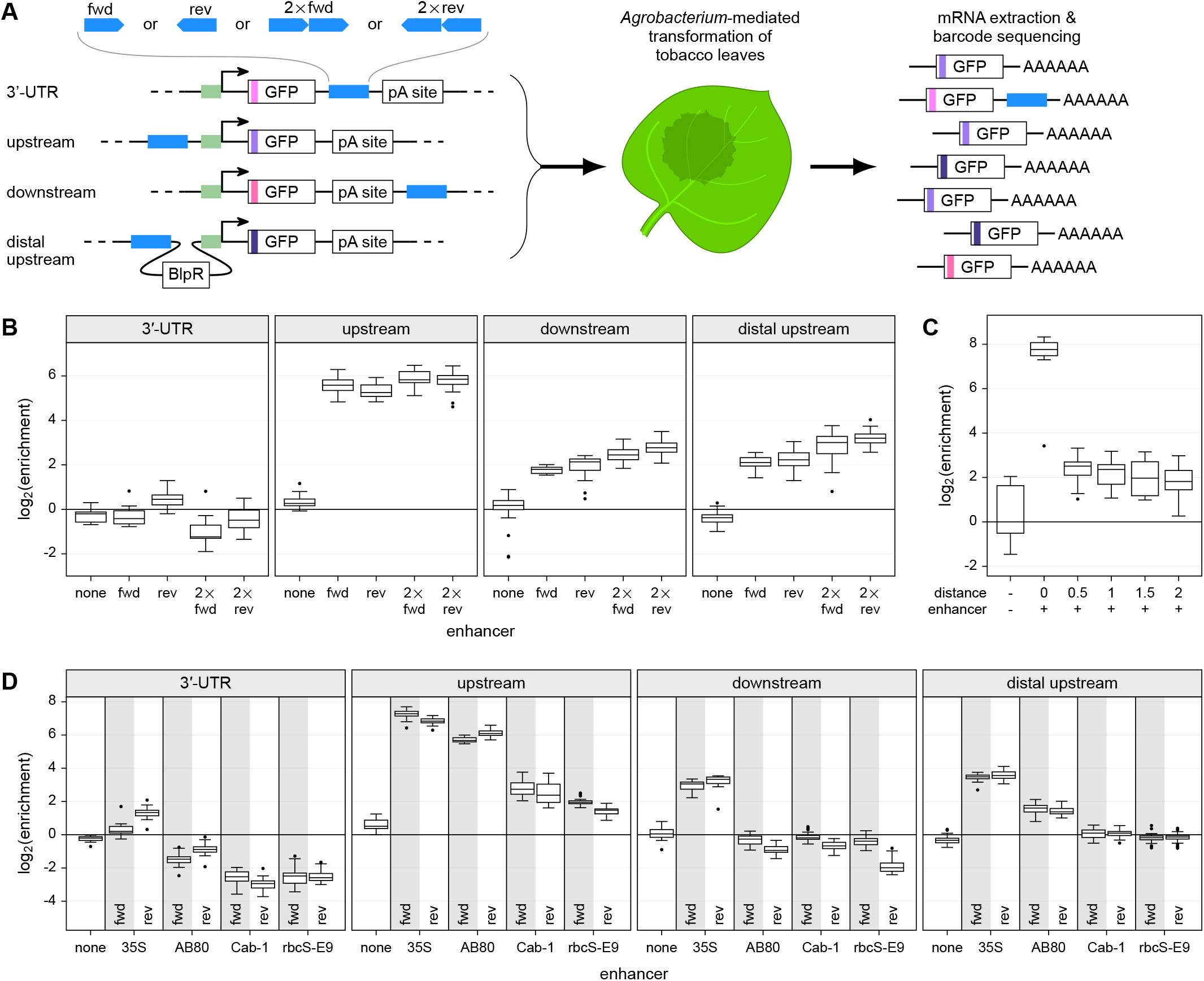
The positioning of enhancers has a pronounced impact on their activity. **(A)** Scheme of the tobacco STARR-seq assay. All constructs are driven by the 35S minimal promoter (green). Enhancers (blue) are inserted in the indicated orientation and position. Barcodes (shades of purple) are inserted in the GFP open reading frame. BlpR, phosphinothricin resistance gene; pA site, poly-adenylation site. **(B)** STARR-seq was performed with constructs harboring a single or double (2×) 35S enhancer in the indicated positions and orientations. Plots show log_2_(enrichment) of recovered RNA barcodes compared to DNA input. Each boxplot (center line, median; box limits, upper and lower quartiles; whiskers, 1.5x interquartile range; points, outliers) represents all barcodes from three independent replicates combined, as do subsequent ones. **(C)** After introduction of a 2 kb spacer upstream of the minimal promoter, the 35S enhancer was inserted at the indicated distance upstream of the minimal promoter, and STARR-seq was performed. **(D)** The 35S and three known plant enhancers were introduced in either the forward (fwd) or reverse (rev) orientation at the indicated positions and the STARR-seq assay was performed.

To ensure a wide dynamic range of the assay, we systematically analyzed the position- and orientation-dependency of the 35S enhancer (Figure 1A). We used a more generalized version of STARR-seq in which we placed a barcode in the GFP open reading frame. This barcode is linked to the corresponding enhancer variant by next generation sequencing and serves as a readout for the activity of the variant. For each variant, we used 5-10 constructs with different barcodes. This barcode redundancy helps to mitigate potential effects that an individual barcode might have on the transcript level. As expected, the 35S enhancer was active in either orientation and both up- and downstream of the reporter gene (Figure 1B). Similar to previous observations (Fang et al., 1989), the activity of the 35S enhancer was lower when present downstream of the gene as compared to upstream of the minimal promoter. In contrast to the mammalian system, when placed in the 3′-UTR, the enhancer had almost no activity. Addition of a second copy of the enhancer in the ‘downstream’ and ‘distal upstream’ positions led on average to a 70% increase in transcript levels as compared to a single enhancer, while a second copy in the ‘upstream’ position increased transcript levels by only 30% (Figure 1B). These observations suggest that the transcriptional activation caused by a single 35S enhancer directly upstream of the minimal promoter is already close to the maximum level detectable in our assay.

We observed the strongest activation of transcription with the enhancer immediately upstream of the minimal 35S promoter, and lower levels when the enhancer was placed about 1.5 kb away from the promoter as in the ‘downstream’ and ‘distal upstream’ constructs (Figures 1A and 1B). To characterize the distance-activity relationship, we inserted the 35S enhancer at different positions within a 2 kb spacer upstream of the minimal promoter (Figure 1C). Enhancer activity was strongest immediately upstream of the promoter. However, enhancer activity was greatly reduced by 500 bp or more of spacer between the enhancer and promoter (Figure 1C), consistent with a previously described distance-dependent decrease of 35S enhancer activity (Odell et al., 1988).

To test if the observed position-dependency is unique to the 35S enhancer, we assayed three additional enhancers derived from the *Pisum sativum* AB80 and rbcS-E9 genes and the wheat Cab-1 gene (Simpson et al., 1986; Fluhr et al., 1986; Nagy et al., 1987; Giuliano et al., 1988; Fejes et al., 1990; Argüello et al., 1992; Gotor et al., 1993). Similar to the 35S enhancer, these enhancers were orientation-independent and most active immediately upstream of the promoter, and they did not activate transcription when placed in the 3′-UTR (Figure 1D).

### The 35S enhancer is not active in the transcribed region

Although previous STARR-seq studies placed candidate enhancer fragments in the 3′-UTR (Arnold et al., 2013; Sun et al., 2019; Ricci et al., 2019), enhancers in this position were not active in our system. To test if the lack of enhancer activity in the 3′-UTR is specific to our assay in transiently transformed tobacco leaves or a more general feature of enhancers in plants, we performed STARR-seq in maize (*Zea mays* L. cultivar B73) protoplasts (Figure 2A). The results with maize protoplasts were qualitatively similar to those from the assay in tobacco leaves. The 35S enhancer was most active upstream of the minimal promoter, and its activity was greatly reduced when placed in the 3′-UTR (Figure 2B). Quantitatively, the activity of the 35S enhancer in the upstream position was lower in the maize protoplasts compared to that observed in tobacco leaves. However, the activity of the 35S enhancer in the 3′-UTR position was slightly higher in maize protoplasts than in tobacco leaves (compare Figures 1B and 2B).

**Figure 2.**
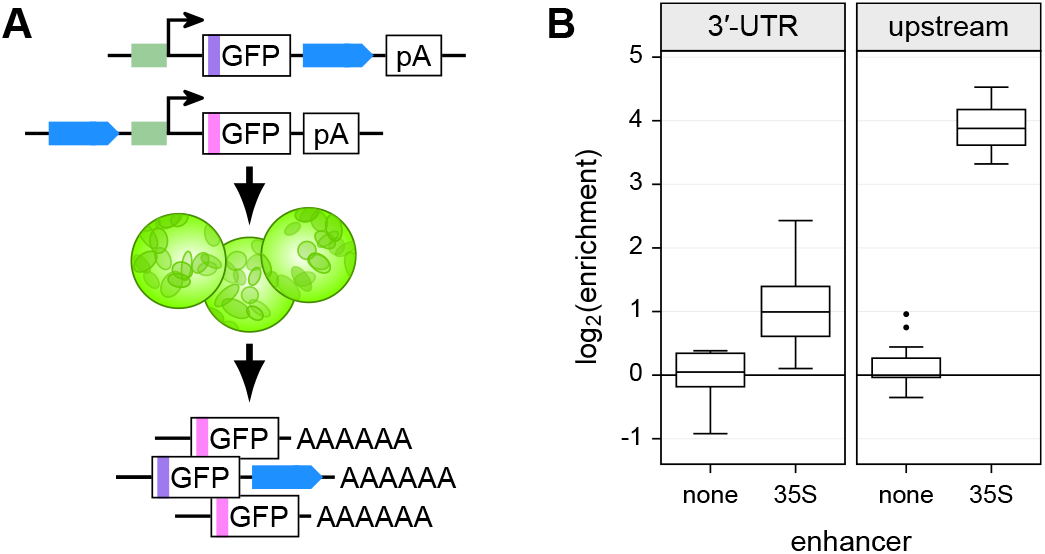
The 35S enhancer is more active upstream of the minimal promoter in Maize protoplasts. **(A)** Scheme of the Maize protoplast STARR-seq assay. **(B)** Maize protoplasts were transformed with constructs without an enhancer (none) or with the 35S enhancer (35S) in the indicated position and subjected to the STARR-seq assay.

To explain the low activity of the 35S enhancer in the 3′-UTR, we hypothesized that such an mRNA could be degraded by nonsense-mediated decay, as long 3′-UTRs can subject mRNAs to this decay pathway (Kertész et al., 2006). To test whether the 35S enhancer in the 3′-UTR destabilizes the mRNA by promoting nonsense-mediated decay, we inserted the unstructured region from the Turnip crinkle virus 3′-UTR, shown to reduce nonsense-mediated decay (May et al., 2018), in between the stop codon and the enhancer. However, insertion of this region further reduced transcript levels when the 35S enhancer was placed in the 3′-UTR (Figures 3A and 3B). We next asked whether insertion of the 35S enhancer in an intron, which would also be transcribed, could confer transcriptional activation, but found that it did not (Figures 3A and 3C). Furthermore, combining an upstream AB80 enhancer with a 35S enhancer within the 3′-UTR transcribed region considerably reduced transcription compared to that from the AB80 enhancer alone (Figure 3D). Taken together, these findings demonstrate that the 35S enhancer residing within the transcribed region is not active in our system. Therefore, for subsequent experiments, we placed the enhancer fragments directly upstream of the minimal promoter, barcoding the reporter amplicons to enable detection by RNA-seq. A similar approach with a barcode in the transcript was used in previous studies of enhancers in human cells (Kwasnieski et al., 2012; Inoue et al., 2019).

**Figure 3.**
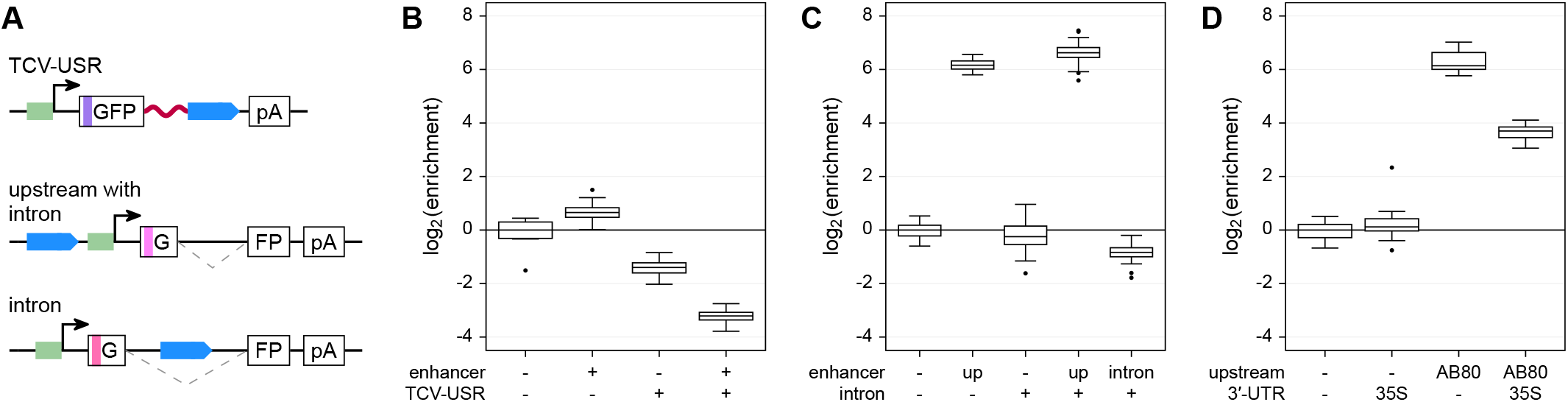
The 35S enhancer is not active in the transcribed region. **(A)** Scheme of the constructs used in this figure. All constructs contain a 35S minimal promoter (green) controlling expression of a GFP reporter gene with or without an intron. In some constructs, an unstructured region of the Turnip crinkle virus 3′-UTR (TCV-USR) was inserted after the stop codon. The 35S core enhancer (blue) was inserted into the indicated positions. **(B)** Constructs with the TCV-USR inserted after the stop codon were subjected to the STARR-seq assay. **(C)** STARR-seq with constructs harboring an intron in the GFP open reading frame. The 35S enhancer was inserted upstream (up) of the promoter or into the intron (intron). **(D)** STARR-seq using constructs with the AB80 enhancer upstream of the minimal promoter and/or the 35S enhancer in the 3′-UTR.

### The tobacco STARR-seq assay can detect enhancer fragments and their light-dependency

The AB80, Cab-1 and rbcS-E9 enhancers are activated by light (Simpson et al., 1986; Nagy et al., 1987; Fluhr et al., 1986). We tested the light-dependency of these enhancers in our assay system by placing the transformed plants in the dark prior to mRNA extraction. The AB80 and Cab-1 enhancers demonstrated decreased activity in the dark. Although the activity of the rbcS-E9 enhancer also showed a response to light, in this case the activity was higher in the dark (Figure 4). A previous study found higher expression of *Arabidopsis thaliana rbcS* genes in extracts from dark-grown plant cells compared to those from light-grown ones, with reversal of this tendency upon reconstitution of chromatin (Ido et al., 2016). It is not clear if plant cells deposit nucleosomes onto the T-DNA harboring the reporter construct, but even if they do, nucleosome positioning and modifications might differ from those found at the endogenous loci of the enhancers.

**Figure 4.**
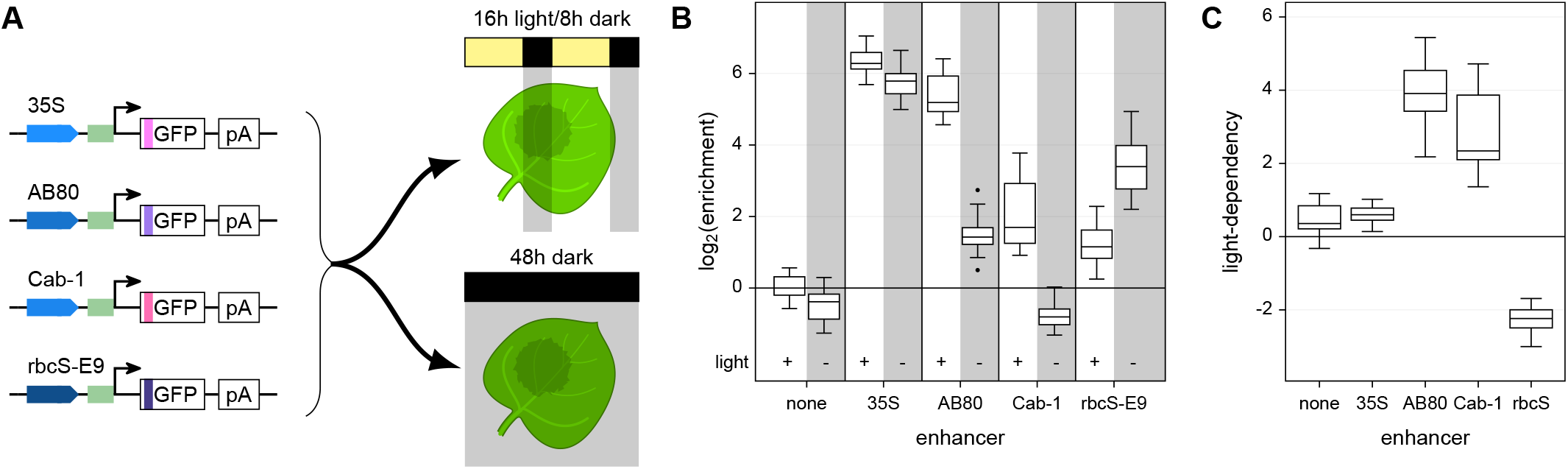
STARR-seq can detect light-dependency of plant enhancers. **(A)** Tobacco leaves were infiltrated with reporter constructs driven by the indicated enhancers. The plants were then grown for two days in normal light/dark cycles or completely in the dark prior to mRNA extraction. **(B)** The activity of the indicated enhancers was determined after growth in normal light/dark cycles (+ light) or in the dark (− light). **(C)** Light-dependency (log_2_(enrichment^light^/enrichment^dark^) was determined for the indicated enhancers.

Next, we tested if the assay could detect enhancer signatures among randomly fragmented DNA sequences from a plasmid containing embedded enhancers. We constructed a plasmid harboring the 35S, AB80, Cab-1, and rbcS-E9 enhancers. We fragmented the plasmid using Tn5 transposase and inserted the fragments upstream of the 35S minimal promoter to generate a fragment library for use in the STARR-seq assay (Figure 5A). This fragment library consisted of approximately 6,200 fragments linked to a total of ~50,000 barcodes. About 40,000 (80%) of these barcodes were recovered with at least 5 counts from the extracted mRNAs. The STARR-seq assay identified the known enhancers as the regions with highest enrichment values (Figure 5B). As expected, the orientation in which the fragments were cloned into the STARR-seq plasmid did not affect their enrichment (Supplemental Figure 1A). This result confirms that the fragments act as enhancers instead of as autonomous promoters, whose activity would be orientation-dependent. The assay was highly reproducible, with good correlation across replicates for the individual barcodes (Spearman’s *ρ* = 0.79–0.82, Supplemental Figure 1B). The correlation further improved if the enrichment of all barcodes linked to the same fragment was aggregated (Spearman’s *ρ*= 0.80–0.85, Supplemental Figure 1C). Replicate correlations were similar for all STARR-seq experiments in this study (Spearman’s *ρ* ≥ 0.6 for barcodes and ≥ 0.7 for fragments or variants, Supplemental Table 1).

**Figure 5.**
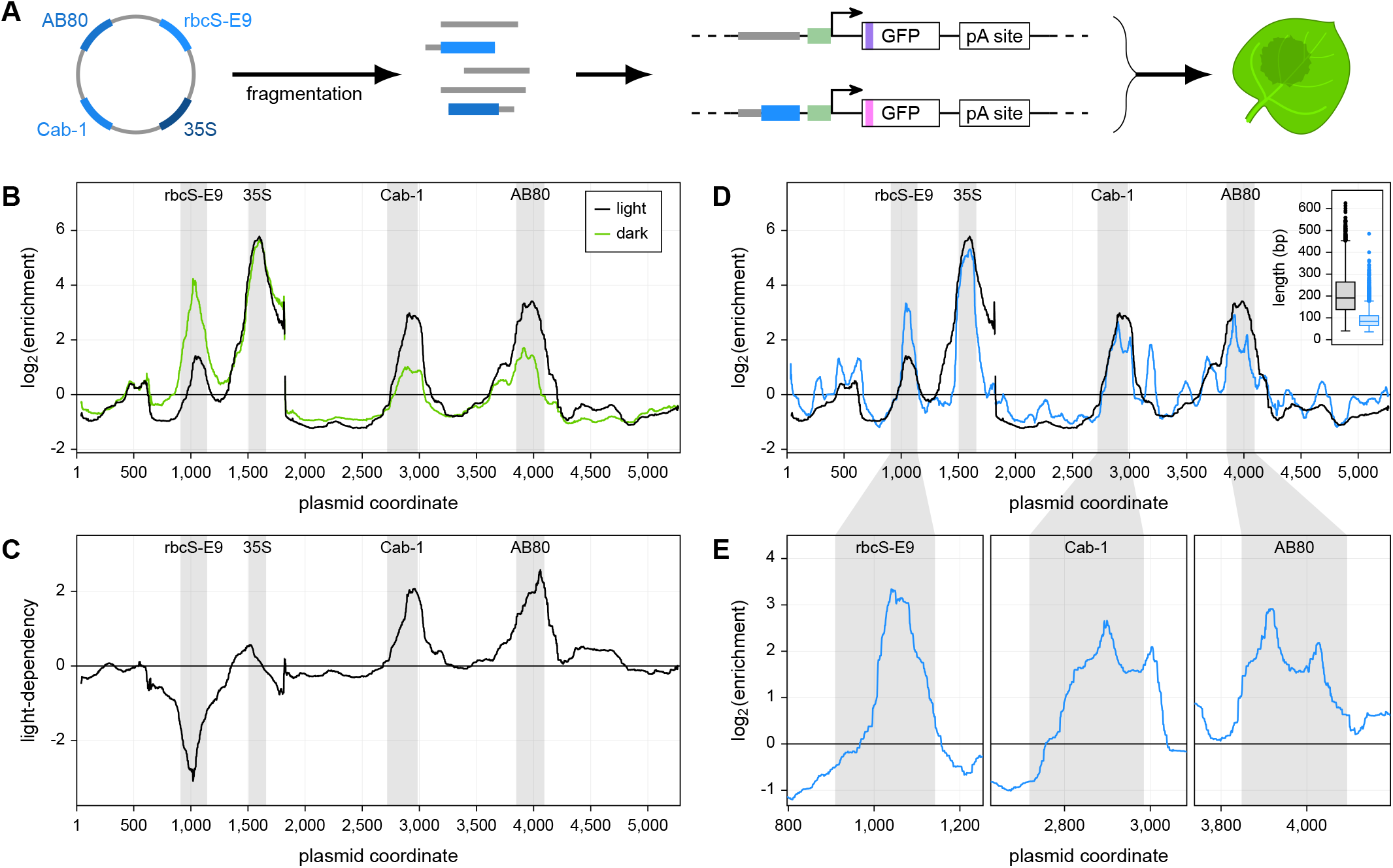
The tobacco STARR-seq assay identifies condition-specific enhancers fragments. **(A)** A plasmid harboring the indicated enhancers was fragmented. The fragments were inserted in the upstream position of the STARR-seq construct and their activity was measured by the STARR-seq assay. **(B)** Plants were grown for two days in normal light/dark cycles (light, black line) or completely in the dark (dark, blue line) prior to mRNA extraction. The log_2_(enrichment) of RNA expression over input of all fragments at each position was averaged. **(C)** Light-dependency (log_2_(enrichment^light^/enrichment^dark^) was determined for each base of the original plasmid. **(D)** The STARR-seq assay was performed with plasmid fragment libraries with different fragment length distributions (see inset), and log_2_(enrichment) for each fragment library is shown across the whole plasmid. **(E)** log_2_(enrichment) obtained from the library with shorter fragments is shown in more detail for regions of interest. Positions in the original plasmid that contain enhancers are shaded in gray.

We also used the fragment library in a STARR-seq experiment with plants kept in the dark prior to mRNA extraction to test for light-dependency. We observed the expected changes in enrichment (Figure 4), with the AB80 and Cab-1 enhancers less active and the rbcS-E9 enhancer more active in the light-deprived plants (Figures 5B and 5C). We conclude that the STARR-seq assay established in this study can identify enhancers in a condition-specific manner.

### The tobacco STARR-seq assay can pinpoint functional enhancer elements

To further reveal individual elements of enhancers, we repeated the screen with a second library (5,700 fragments with a total of 73,000 barcodes, more than 95% of which were recovered from the mRNA) that contained shorter fragments (median length 84 bp vs. 191 bp in the initial library, Figure 5D). As these shorter fragments were, on average, well below the size of the full-length enhancers, they are unlikely to contain all the elements required for maximum activity. The shorter fragments split the peaks of the AB80 and Cab-1 enhancers into two subpeaks, suggesting that these enhancers contain at least two independent functional elements. The sole functional element of the rbcS-E9 enhancer resided in the 3′ half of the tested region (Figures 5D and 5E).

Having established the capacity of the assay to distinguish enhancer subdomains, we tested its suitability for conducting saturation mutagenesis of *cis*-regulatory elements. To do so, we array-synthesized all possible single nucleotide substitution, deletion, and insertion variants of the minimal promoter and of the 35S enhancer as two separate variant pools, and subjected the two pools to STARR-seq. Approximately 98% of all possible variants were linked to at least one barcode in the input library, and mRNAs corresponding to over 99% of these were recovered from the tobacco leaves. We first assayed the activity of variants of a 46 bp region containing the 35S minimal promoter, in constructs with and without an enhancer. The effects of the individual mutations were similar in both contexts (Supplemental Fig. 2). As expected, mutations that disrupt the TATA box (positions 16-22) had a strong negative impact on promoter activity, while most others had a weak effect or no effect (Figures 6A and 6B, Supplemental Data Set 1).

**Figure 6.**
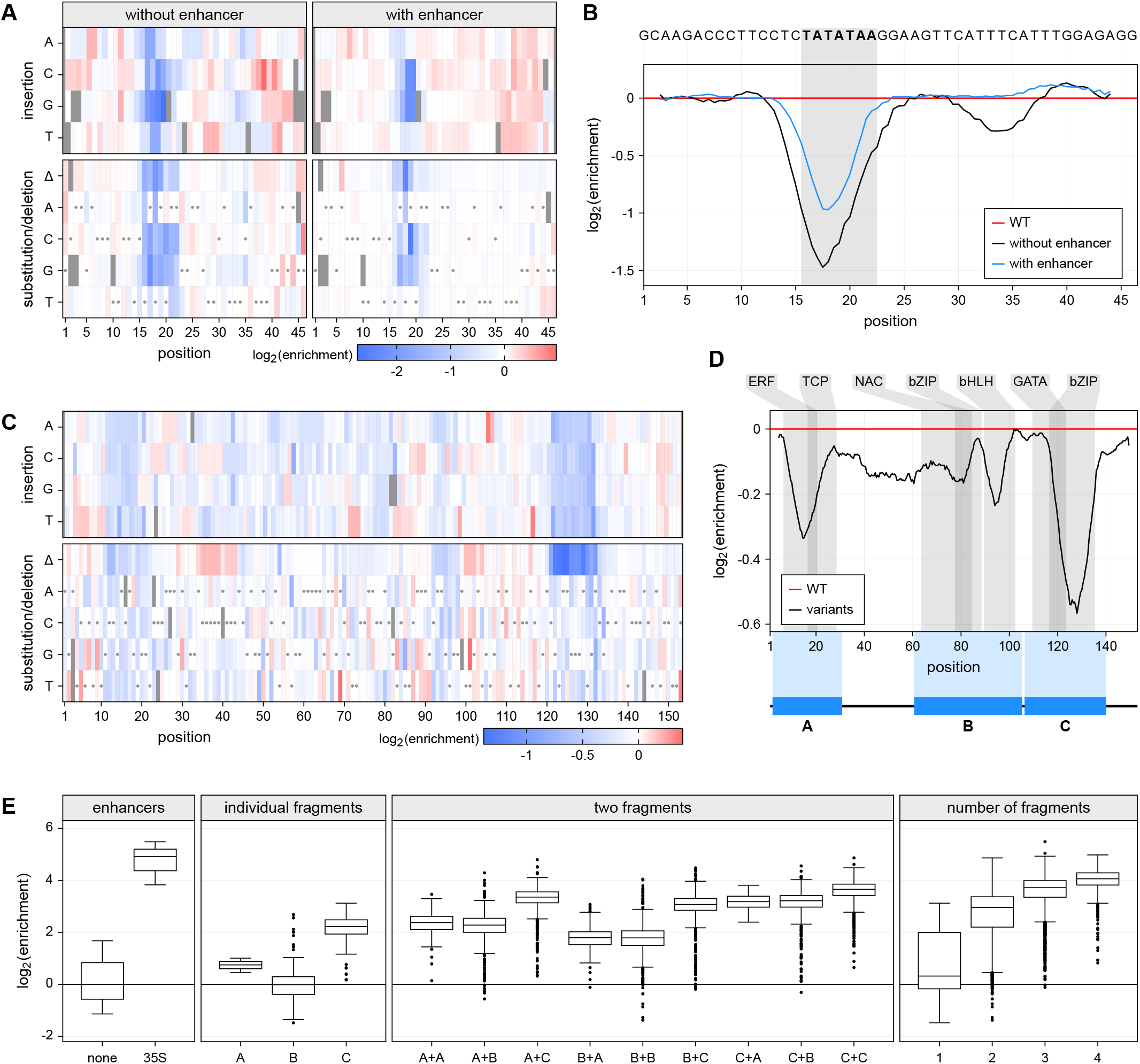
Saturation mutagenesis identifies functional elements in the 35S promoter and enhancer. **(A)** All possible single nucleotide variants of the 35S minimal promoter were inserted into constructs with or without the 35S enhancer. The enrichment of the individual promoter insertions, substitutions and deletions was measured by the STARR-seq assay, normalized to the wild type variant, and plotted as a heatmap. Missing values are shown in black and wild type variants are marked with gray dot. **(B)** A sliding average (window size = 4 bp) of the positional mean enrichment scores for all substitutions, insertions and deletions was determined. The TATA box is highlighted in gray. **(C)** The enrichment of all possible single nucleotide insertions, substitutions and deletions of the 35S core enhancer was determined as in **(A)**. **(D)** A sliding average (window size = 8 bp) of the positional mean enrichment was determined via the STARR-seq assay. Predicted binding sites for the transcription factors from the indicated families are highlighted in gray. **(E)** Three fragments (A, B, C; see **D**) of the 35S enhancer were inserted into the STARR-seq plasmid in random number and order, and assayed for their enhancer activity.

In contrast to the minimal promoter, the 35S enhancer contained several mutation-sensitive regions (Figures 6C and 6D, Supplemental Data Set 2). These regions co-localize with predicted transcription factor binding sites (Tian et al., 2020; Grant et al., 2011). Mutations in positions 116-135 were especially deleterious. This region, previously implicated in enhancer activity, can be bound by the tobacco activation sequence factor 1 (ASF-1), a complex containing the bZIP transcription factor TGA2.2 (Fang et al., 1989; Benfey et al., 1990; Lam et al., 1989; Niggeweg et al., 2000). Similarly, we observed mutational sensitivity of the 35S enhancer in positions 95-115, which contain a binding site for the bHLH transcription factor complex ASF-2 (Lam and Chua, 1989). A third mutation-sensitive region in positions 7-28 is predicted to be bound by ERF and TCP transcription factors.

### Enhancer fragments can be combined to build synthetic enhancers

To demonstrate that these mutation-sensitive regions possess enhancer activity, we split the enhancer into three fragments that span positions 1-30 (A), 60-105 (B), and 106-140 (C) (Figure 6D). These fragments were cloned in one to four copies on average, in random order, and the enhancer activity of the resulting constructs was determined. We identified 100 different constructs linked to a total of 29,000 barcodes, 95% of which were present in the extracted mRNAs. Fragments A and C alone were sufficient to activate transcription, while fragment B was active only in the presence of a second fragment (Figure 6E). In line with our observations from the enhancer mutagenesis, fragment C had the highest activity. The greater the number of fragments in a construct, the higher its activity. However, even four fragments combined did not reach the level of transcription achieved with the full-length enhancer, indicating that the sequences excluded from the A, B and C fragments contribute to enhancer activity, either directly or by providing the correct spacing for the fragments (Figure 6E). Although spacing may play a role in enhancer activity, the order of the fragments had only weak effects (Supplemental Figure 3). Taken together, we demonstrate that this assay can identify functional enhancer elements that can be recombined to create synthetic enhancers of varying strength.

## DISCUSSION

In this study, we developed a massively parallel reporter assay in tobacco plants that can identify DNA regions with enhancer or promoter activity and can dissect these regions to characterize functional sequences with single nucleotide resolution. The assay does not depend on efficient protoplasting and transformation protocols, which have been established only for a limited number of species and tissues. Furthermore, in contrast to protoplasts, the *in planta* system is more robust and can be exposed to a variety of environmental conditions to detect condition-specific *cis*-regulatory elements. Indeed, our tobacco STARR-seq assay can detect enhancer light-dependency. Such condition-specific *cis*-regulatory elements could play important roles in future genetic engineering efforts to help plants adapt to a rapidly changing environment.

We observed in our experiments that the tested enhancers were not active when placed in the transcribed region. Other studies have shown that plant genes can contain elements in their transcribed region, especially in the first intron, that drastically increase their expression (Callis et al., 1987; Rose and Last, 1997; Rose, 2004; Samadder et al., 2008; Laxa et al., 2016; Laxa, 2017). However, the increased expression levels could have been the result of enhanced transcription or translation, improved mRNA processing, export, or stability, or a combination of these mechanisms. Few studies have dissected these potential mechanisms, and these have generally found that enhanced transcription played no role, or only a relatively small role, in the overall expression increase (Rose and Last, 1997; Samadder et al., 2008; Laxa et al., 2016). The apparent absence of strong transcriptional enhancers in the transcribed region of plant genes could be due to any of several reasons. The constraints placed on such regions to enable efficient mRNA processing and translation might not be compatible with the requirements for enhancers. Alternatively, strong binding of transcription factors within the transcribed region could inhibit transcription by physically blocking the RNA polymerase. Future studies will be required to address this issue in plants.

Comparing the activity of the 35S enhancer in transiently transformed tobacco leaves to its activity in maize protoplasts, a general trend of high activity upstream of the minimal promoter and low activity within the 3′-UTR was observed, but the levels differed between the two systems. Previous studies have reported that the 35S promoter constructs encompassing the 35S minimal promoter and enhancer are more active in dicots like tobacco than monocots such as maize and rice (Christensen et al., 1992; Bruce et al., 1989). In agreement with these studies, we detected higher activity of the 35S enhancer upstream of the minimal promoter in tobacco compared to maize. In contrast, the maize system led to more 35S enhancer activity than the tobacco system when the enhancer was inserted into the 3′-UTR, a possible effect of species-specific differences in the tolerance of an enhancer within the transcribed region. Consistent with these species differences, effects of intron-mediated enhancement of gene expression are stronger in monocots than dicots (Samadder et al., 2008). Alternatively, the physical state of the reporter construct-containing DNA could influence enhancer activity. In maize protoplasts, the reporter is expressed from a supercoiled plasmid, whereas it resides on linear T-DNA molecules in the transiently transformed tobacco cells. Linear and supercoiled DNA probably display different looping behavior that could influence medium-to long-range enhancer-promoter interactions.

In the two previous plant STARR-seq studies (Sun et al., 2019; Ricci et al., 2019), the enhancer candidates were placed in the 3′-UTR of the reporter gene. While these studies successfully identified several strong enhancers, our results indicate that the dynamic range in these studies might have been improved by altering the placement of the enhancer. The reporter design we used here with enhancer candidates immediately upstream of the minimal promoter yields a high signal-to-noise ratio and enables confident discovery of intermediate and weak enhancers. This design requires an additional sequencing step to link the enhancer candidates to the corresponding barcodes. However, the barcodes are short and of the same length, whereas the length of enhancer candidates can vary considerably. Cloning these highly variable enhancer sequences into the 3′-UTR may have a profound impact on the stability of the resulting mRNAs and how readily they can be recovered and sequenced.

Apart from the enhancer placement, the plant species (tobacco, rice or maize) and the recipient cells/tissue (protoplasts or intact leaves) differ between this study and the two previous plant STARR-seq studies (Sun et al., 2019; Ricci et al., 2019). As 35S enhancer activity in tobacco leaves and maize protoplasts showed similar trends, the choice of a model system for future STARR-seq studies will likely depend on what mode of transformation is most efficient in the target species. While several species are amenable to transient transformation with *Agrobacteria* (Wroblewski et al., 2005; Andrieu et al., 2012; Zheng et al., 2012; Bond et al., 2016), others might be more easily transformed as protoplasts (Sheen, 1990; Zhang et al., 2011; Nanjareddy et al., 2016). Transformation efficiency will pose a limit to the maximum library size that can be screened. The two previous plant STARR-seq studies transformed 15-30 million protoplasts (Sun et al., 2019; Ricci et al., 2019). The largest library in this study contained about 73,000 barcodes, of which we detected more than 95% in the extracted mRNAs. In experiments with a larger library, we could recover 250,000 fragments from a single tobacco leaf. As the extraction of mRNAs from 100 tobacco leaves can be performed in a single day, library sizes similar to the ones used in the other plant STARR-seq studies should be feasible.

Due to the widespread conservation of transcription factors in the plant lineage (Lehti-Shiu et al., 2017; Wilhelmsson et al., 2017), the enhancer elements identified in tobacco leaves will likely be active in many other plant species. Furthermore, the STARR-seq assay described herein can potentially be performed in any species or tissue that can be transiently transformed by *Agrobacteria*. Apart from enhancers and promoters, the assay can likely be adapted to screen for silencers and insulators — *cis*-regulatory elements that are known from animals but have, so far, not been detected in plants.

Taken together, we describe a plant STARR-seq assay that is applicable to enhancer screens for any plant species to analyze plant gene regulation and to identify promising building blocks for future genetic engineering efforts. The data generated by these screens and subsequent saturation mutagenesis will enable deep learning approaches to identify defining characteristics of plant enhancers.

## METHODS

### Plasmid construction and library creation

The STARR-seq plasmids used herein are based on the pGreen plasmid (Hellens et al., 2000). In their T-DNA region, they harbor a phosphinothricin resistance gene (BlpR) and the GFP reporter construct terminated by the polyA site of the *Arabidopsis thaliana* ribulose bisphosphate carboxylase small chain 1A gene. These plasmids were deposited at Addgene (Addgene #149416-149422, https://www.addgene.org/). The 35S minimal promoter followed by the synthetic 5′-UTR synJ (Kanoria and Burma, 2012) (ACACGCTGGAATTCTAGTATACTAAACC), an ATG start codon and a 15 bp random barcode (VNNVNNVNNVNNVNN) was cloned in front of the second codon of GFP by Golden Gate cloning (Engler et al., 2008). All primers are listed in Supplemental Data Set 3. Enhancers or DNA fragments were inserted by Golden Gate cloning into the indicated positions. The 2 kb spacer used in Figure 1C was derived from enCas9 in pEvolvR-enCas9-PolI3M-TBD (Halperin et al., 2018) (Addgene #113077, https://www.addgene.org/113077/). For constructs with full-length enhancers (Figures 1B to 1D, 2, 3, and 4), 5-10 uniquely barcoded variants were used. The sequences of the 35S, AB80, Cab-1, and rbcS-E9 enhancers are listed in Supplemental Table 2. These enhancers were inserted into the SacI, XbaI, XhoI, and SfoI sites of the plasmid pZS*11-yfp0 (Subramaniam et al., 2013) (Addgene #53241, https://www.addgene.org/53241/). The resulting plasmid (pZS*11_4enh, Addgene #149423, https://www.addgene.org/149423/) was fragmented with Nextera Tn5 transposase (Illumina) and the fragments were amplified with primers containing adapters suitable for Golden Gate cloning. The single nucleotide variants of the 35S promoter and enhancer were ordered as an oligonucleotide array from Twist Bioscience (Figures 6A to 6D). The libraries were bottlenecked to approximately 10 barcodes per variant. Fragments A, B, and C (Figure 6D) were ordered as oligonucleotide with 5′ GTGATG overhangs, mixed with Golden Gate cloning adaptors and ligated with T4 DNA ligase. The IV2 intron was inserted into GFP at position 103. The unstructured region of the Turnip crinkle virus 3′-UTR (TACGGTAATAGTGTAGTCTTCTCATCTTAGTAGTTAGCTCTCTCTTATATT) was inserted after the GFP stop codon. Next generation sequencing on an Illumina NextSeq platform was used to link the inserted fragments and barcodes. The STARR-seq plasmid libraries were introduced into *Agrobacterium tumefaciens* GV3101 strain harboring the helper plasmid pSoup (Hellens et al., 2000) by electroporation.

### Plant cultivation and transformation

Tobacco (*Nicotiana benthamiana*) was grown in soil (Sunshine Mix #4) at 25 °C in a long-day photoperiod (16 h light and 8 h dark; fluorescent light intensity 300 μmol/m^2^/s). Plants were transformed 3 - 4 weeks after germination. For transformation, an overnight culture of *Agrobacterium tumefaciens* was diluted into 50 mL YEP medium and grown at 28 °C to an OD of approximately 1. A 5 mL input sample of the cells was taken, and plasmids were isolated from it. The remaining cells were harvested and resuspended in 50 mL induction medium (M9 medium supplemented with 1% glucose, 10 mM MES, pH 5.2, 100 μM CaCl_2_, 2 mM MgSO_4_, and 100 μM acetosyringone). After overnight growth, the bacteria were harvested, resuspended in infiltration solution (10 mM MES, pH 5.2, 10 mM MgCl_2_, 150 μM acetosyringone, and 5 μM lipoic acid) to an OD of 1 and infiltrated into the first two mature leaves of 2 - 4 tobacco plants. The plants were further grown for 48 h under normal conditions or in the dark prior to mRNA extraction.

### Maize protoplast generation and transformation

We used a slightly modified version of a published protoplasting and electroporation protocol (Sheen, 1990). Maize (*Zea mays* L. cultivar B73) seeds were germinated for 4 days in the light and the seedlings were grown in soil at 25 °C in the dark for 9 days. The center 8-10 cm of the second leaf from 7-9 plants were cut into thin strips perpendicular to the veins and immediately submerged in 10 mL protoplasting solution (0.6 M mannitol, 10 mM MES, 15 mg/mL cellulase R-10 [GoldBio], 3 mg/mL Macerozyme R-10 [GoldBio], 1 mM CaCl_2_, 5 mM β-mercaptoethanol, pH 5.7). The mixture was covered to keep out light, vacuum infiltrated for 30 min, and incubated with 40 rpm shaking for 2 h. Protoplasts were released with 80 rpm shaking for 5 min and filtered through a 40 μm filter. The protoplasts were harvested by centrifugation (3 min at 200 x g) in a round bottom glass tube and washed with 3 mL electroporation solution (0.6 M mannitol, 4 mM MES, 20 mM KCl, pH 5.7). After centrifugation (2 min at 200 x g), the cells were resuspended in 3 mL ice cold electroporation solution and counted. Approximately 1 million cells were mixed with 25 μg plasmid DNA in a total volume of 300 μL, transferred to a 4 mm electroporation cuvette and incubated for 5 min on ice. The cells were electroporated (300 V, 25 μFD, 400 Ω) and 900 μL incubation buffer (0.6 M mannitol, 4 mM MES, 4 mM KCL, pH 5.7) was added. After 10 min incubation on ice, the cells were further diluted with 1.2 mL incubation buffer and kept at 25 °C in the dark for 16 h before mRNA collection.

### STARR-seq assay

For each STARR-seq experiment with tobacco plants at least 3 independent biological replicates were performed. Different plants and fresh *Agrobacterium* cultures were used for each biological replicate and the replicates were performed on different days. Depending on library size, two samples of 2–3 leaves were collected from a total of 2–4 plants. They were frozen in liquid nitrogen, ground in a mortar and immediately resuspended in 5 mL TRIzol (Thermo Fisher Scientific). The suspension was cleared by centrifugation (5 min at 4,000 x g, 4 °C), and the supernatant was thoroughly mixed with 2 mL chloroform. After centrifugation (15 min at 4,000 x g, 4 °C), the upper, aqueous phase was transferred to a new tube, mixed with 1 mL chloroform and centrifuged again (15 min at 4,000 x g, 4 °C). 2.4 mL of the upper, aqueous phase were transferred to new tubes, and RNA was precipitated with 240 μL 8 M LiCl and 6 mL 100% ethanol by incubation at −80 °C for 15 min. The RNA was pelleted (30 min at 4,000 x g, 4 °C), washed with 2 mL 70% ethanol, centrifuged again (5 min at 4,000 x g, 4 °C), and resuspended in 500 μL nuclease-free water. mRNAs were isolated from this solution using 100 μL magnetic Oligo(dT)_25_ beads (Thermo Fisher Scientific) according to the manufacturer’s protocol. The mRNAs were eluted in 40 μL. The two samples per library were pooled and supplemented with DNase I buffer, 10 mM MnCl_2_, 2 μL DNase I (Thermo Fisher Scientific), and 1 μL RNaseOUT (Thermo Fisher Scientific). After 1 h incubation at 37 °C, 2 μL 20 mg/mL glycogen (Thermo Fisher Scientific), 10 μL 8 M LiCl, and 250 μL 100% ethanol were added to the samples. Following precipitation at −80 °C, centrifugation (30 min at 20,000 x g, 4 °C), and washing with 200 μL 70% ethanol (5 min at 20,000 x g, 4 °C), the pellet was resuspended in 100 μL nuclease-free water. Eight reactions with 5 μL mRNA each and a GFP construct-specific primer (GAACTTGTGGCCGTTTACG) were prepared for cDNA synthesis using SuperScript IV reverse transcriptase (Thermo Fisher Scientific) according to the manufacturer’s instructions. Half of the reactions were used as no reverse transcription control and the enzyme was replaced with water. After cDNA synthesis, the reactions were pooled and purified with DNA Clean & Concentrator-5 columns (Zymo Research). The barcode region was amplified with 10–20 cycles of PCR and read out by next generation sequencing on an Illumina NextSeq platform.

For the STARR-seq assay in maize protoplasts, we performed 3 independent biological replicates on different days with different plants. Transformed protoplasts were harvested by centrifugation (3 min at 200 x g, 4 °C) 16 h after electroporation. The protoplasts were washed three times with 1 mL incubation buffer and centrifuged for 2 min at 200 x g and 4 °C. The cells were resuspended in 300 μL TRIzol (Thermo Fisher Scientific) and incubated for 5 min at room temperature. The suspension was thoroughly mixed with 60 μL chloroform and centrifuged (15 min at 20,000 x g, 4 °C). The upper, aqueous phase was transferred to a new tube, mixed with 60 μL chloroform and centrifuged again (15 min at 20,000 x g, 4 °C). RNA was precipitated from 200 μL of the supernatant with 1 μL 20 mg/mL glycogen (Thermo Fisher Scientific), 20 μL 8 M LiCl, and 600 μL 100% ethanol by incubation at −80 °C for 15 min. After centrifugation (30 min at 20,000 x g, 4 °C), the pellet was washed with 200 μL 70% ethanol, centrifuged again (5 min at 20,000 x g, 4 °C), and resuspended in 200 μL nuclease-free water. mRNAs were isolated from this solution using 50 μL magnetic Oligo(dT)_25_ beads (Thermo Fisher Scientific) according to the manufacturer’s protocol, and the mRNAs were eluted in 40 μL water. DNase I treatment and precipitation were performed as for the mRNAs obtained from tobacco plants but with half the volume. Reverse transcription, purification, PCR amplification and sequencing were performed as for the tobacco samples. For the maize protoplast STARR-seq, the plasmid library used for electroporation was sequenced as the input sample.

### Computational methods

Binding site motifs for *N. benthamiana* transcription factors were obtained from the PlantTFDB (Tian et al., 2020), and FIMO (Grant et al., 2011) was used to predict their occurrence in the 35S core enhancer. Fragments of pZS*11_4enh were aligned to the reference sequence using Bowtie2 (Langmead and Salzberg, 2012). For analysis of the STARR-seq experiments, the reads for each barcode were counted in the input and cDNA samples. Barcode counts below 5 and barcodes present in only one of three replicates were discarded. Barcode enrichment was calculated by dividing the barcode frequency (barcode counts divided by all counts) in the cDNA sample by that in the input sample. For the pZS*11_4enh fragment library (Figure 5 and Supplemental Figure 1) and the mutagenesis (Figures 6A to 6D) experiments, the enrichment of the fragments or variants was calculated as the median enrichment of all barcodes linked to them. Boxplots were created using all corresponding barcodes from all replicates performed and were normalized to the median enrichment of constructs without an enhancer. The enrichment coverage of pZS*11_4enh was calculated by summing up the enrichment of all fragments containing a given nucleotide and dividing this sum by the number of fragments. Nucleotides covered by less than 5 fragments were excluded from analysis. Light-dependency of enhancers or enhancer fragments was calculated as log_2_ of the enrichment in the light condition divided by the enrichment from the dark condition. The code used for analyses is available at https://github.com/tobjores/tobacco-STARR-seq.

## Supporting information

Supplemental Data Set 1

Supplemental Data Set 2

Supplemental Data Set 3

## Accession Numbers

All barcode sequencing reads were deposited in the NCBI Sequence Read Archive under the BioProject accession PRJNA627258 (http://www.ncbi.nlm.nih.gov/bioproject/627258).

## Supplemental Data

**Supplemental Figure 1.** Activity in the STARR-seq assay is insensitive to orientation and reproducible.

**Supplemental Figure 2.** Activity of the promoter variants is correlated with and without an enhancer in the construct.

**Supplemental Figure 3.** The order of 35S enhancer fragments has a subtle influence on enhancer activity.

**Supplemental Table 1.** Replicate correlations for all STARR-seq experiments in this study.

**Supplemental Table 2.** Enhancers sequences used in this study.

**Supplemental Data Set 1.** Enrichment scores for 35S minimal promoter variants.

**Supplemental Data Set 2.** Enrichment scores for 35S core enhancer variants.

**Supplemental Data Set 3.** Primers used in this study.

## ACKNOWLEDGEMENTS

We thank Dr. Cristina M. Alexandre for initial plasmid construction. This work was supported by NSF RESEARCH-PGR 1748843 grant to S.F. and C.Q.

## AUTHOR CONTRIBUTIONS

T.J., J.T., M.W.D., J.T.C., S.F, and C.Q. conceived and interpreted experiments and wrote the manuscript. T.J. and J.T. performed experiments and T.J. conducted data analysis and prepared the figures.

