## Supplemental Figure 1 for "Identification of plant enhancers and their constituent elements by STARR-seq in tobacco leaves"

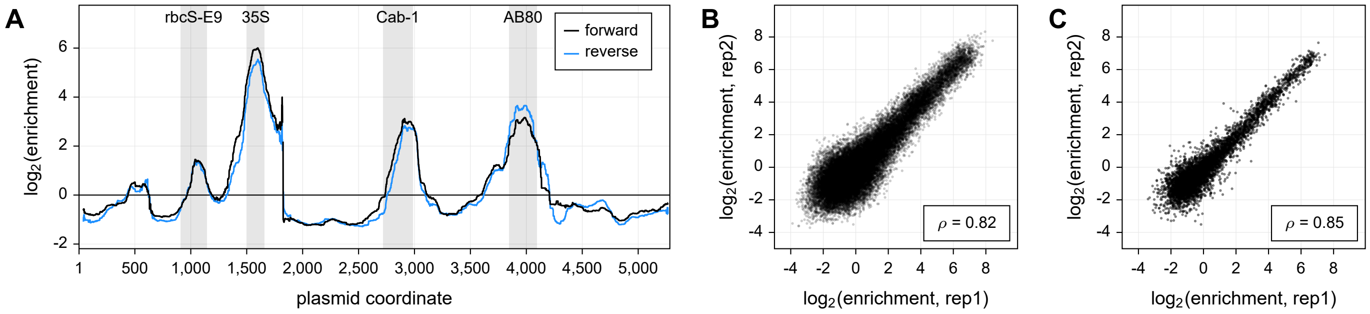


**Supplemental Figure 1.** Activity in the STARR-seq assay is insensitive to orientation and reproducible.

(Supports Figure 5)
**(A)** The data from the STARR-seq assay (see Fig. 5B, light condition) was analyzed separately for fragments inserted in the forward or reverse orientation.
**(B)** Correlation (Spearman's $\rho$) between two (out of three) replicates for individual barcodes.
**(C)** Correlation (Spearman's $\rho$) between two (out of three) replicates for fragments (median enrichment of all linked barcodes).
