## Supplemental Figure 2 for "Identification of plant enhancers and their constituent elements by STARR-seq in tobacco leaves"

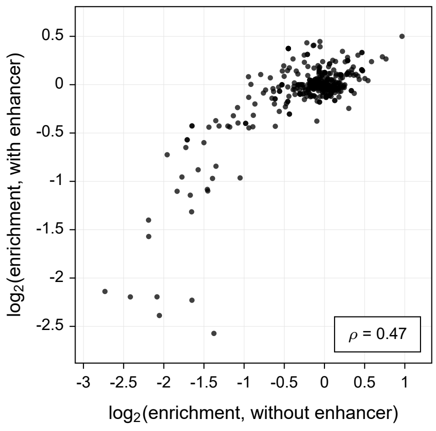


**Supplemental Figure 2.** Activity of the promoter variants is correlated with and without an enhancer in the construct.

(Supports Figure 6)
The enrichment scores for 35S minimal promoter variants (see Fig. 6A) in constructs with or without an upstream 35S enhancer was compared. The correlation (Spearman's $\rho$) is indicated.
