## Supplemental Figure 3 for "Identification of plant enhancers and their constituent elements by STARR-seq in tobacco leaves"

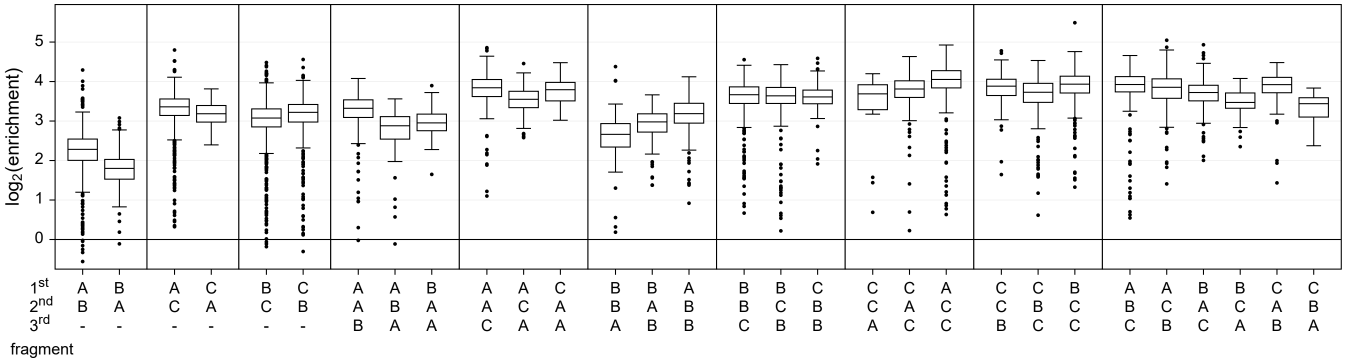


**Supplemental Figure 3.** The order of 35S enhancer fragments has a subtle influence on enhancer activity.

(Supports Figure 6)

Three fragments (A, B, C) of the 35S enhancer (see Fig. 6D) were inserted into the STARR-seq plasmid in random number and order, and assayed for their enhancer activity. Each boxplot (center line, median; box limits, upper and lower quartiles; whiskers, 1.5x interquartile range; points, outliers) represents all barcodes from three independent replicates combined.
