## Supplemental Table 1 for "Identification of plant enhancers and their constituent elements by STARR-seq in tobacco leaves"

**Supplemental Table 1.** Replicate correlations for all STARR-seq experiments in this study.

Spearman's correlation between biologiocal replicates was determined for all STARR-seq experiments. Different plants and fresh Agrobacterium cultures were used for each biological replicate and the replicates were performed on different days.

| Experiment | replicates | | |
| --- | --- | --- | --- |
|  | 1 vs. 2 | 1 vs. 3 | 2 vs. 3 |
| Figure 1B (barcodes) | 0.98 | 0.98 | 1.00 |
| Figure 1C (barcodes) | 0.86 | 0.67 | 0.89 |
| Figure 1D (barcodes) | 0.99 | 0.99 | 0.99 |
| Figure 2B (barcodes) | 0.99 | 0.98 | 0.99 |
| Figure 3B (barcodes) | 0.97 | 0.97 | 0.98 |
| Figure 3C (barcodes) | 1.00 | 0.99 | 1.00 |
| Figure 3D (barcodes) | 1.00 | 1.00 | 0.99 |
| Figure 4B (barcodes, light) | 0.96 | 1.00 | 0.97 |
| Figure 4B (barcodes, dark) | 0.98 | 0.98 | 0.98 |
| Figure 5B (barcodes, light) | 0.82 | 0.79 | 0.80 |
| Figure 5B (fragments, light) | 0.85 | 0.80 | 0.80 |
| Figure 5B (barcodes, dark) | 0.74 | 0.75 | 0.77 |
| Figure 5B (fragments, dark) | 0.71 | 0.73 | 0.74 |
| Figure 5D (barcodes, short fragment library) | 0.67 | 0.70 | 0.65 |
| Figure 5D (fragments, short fragment library) | 0.84 | 0.89 | 0.83 |
| Figure 6A (barcodes) | 0.86 | 0.86 | 0.86 |
| Figure 6A (variants) | 0.91 | 0.91 | 0.90 |
| Figure 6C (barcodes) | 0.63 | 0.65 | 0.60 |
| Figure 6C (variants) | 0.78 | 0.80 | 0.75 |
| Figure 6E (barcodes) | 0.91 | 0.89 | 0.87 |
