## Supplemental Table 2 for "Identification of plant enhancers and their constituent elements by STARR-seq in tobacco leaves"

Supplemental Table 2. Enhancers sequences used in this study.

| Enhancer | Species | Sequence | Ref. |
| --- | --- | --- | --- |
| 35S | Cauliflower mosaic virus | AGATCTCTCTGCCGACAGTGGTCCCAAAGATGGACCCCCACCCACGAGGA  GCATCGTGGAAAAAGAAGACGTTCCAACCACGTCTTCAAAGCAAGTGGAT  TGATGTGACATCTCCACTGACGTAAGGGATGACGCACAATCCCACTATCC  TTC | Fang et al., 1989; Benfey et al., 1990 |
| AB80 | *Pisum sativum* | CTGCCATAATGTCACAATTTTTCTCAAATCTTGTGGCTCTCAAACACTGT  ATAAAACACGACAAATGTGGACCCAAAATATATACCTTACACTTCTGAGT  TAGAGAAGCAGAGCCCCATAATTAAGCCTATTTTATGAAAAAAATAATAT  TATGTTGAGTCATATATCCATAAGAATCCCCACAGTCACACATGGAAGAG  CAGCATTGGATACAAATGATATGAAGATTTTGATCCATGAACGGATT | Simpson et al., 1986 |
| rbcS-E9 | *Pisum sativum* | ACACTAACATCGAATGTACTTATCTCATTAGTTTAAATTATTGTTTGATC  ATGTTTAATCCTTATACTGTTGTTAGTTTTTTCAGTTAGCTTAGTGGGCA  TCTTACACGTGGCATTATTATCCTATTGGTGGCTAATGATAAGGTTAGCA  CACAAAACTTTTCAATCTTGTGTGGTTAATATGACTGCAAAGTTTATCAT  TTTCACAATCCAACAAACTGGTTCTAGGCAGTGG | Fluhr et al., 1986 |
| Cab-1 | *Triticum aestivum* | ATCGAATTTGTTGGGCAAGCGCCCAGTGTGATGTACGTTGGGAAAACTTG  CAAGAGGATGCGACCAAATGAACTGGTAAACCATCCCGTGAGCGTGGCCT  ACACATTTTAAGCCAGCGGACTCTTTCGACTTGTCTTACAAAAGCTGGTC  CAGTCACGAGCCTTAGCCCTAACCATAGCCACAAGTACATCTCATCCATT  TAAGGCCTCTGCGTGCACCAATGGCATCCAAGCTGCAGATTTCTTTTCAC  CACCGTCTCTCTTGTCAG | Nagy et al., 1987 |
